# Virus-induced genome editing in the parasitic plant *Phtheirospermum japonicum*

**DOI:** 10.64898/2026.02.10.705035

**Authors:** Hannes Ruwe, Victoria Zimmer, Thomas Spallek

## Abstract

*Phtheirospermum japonicum* is a genetic model for parasitic Orobanchaceae, a plant family that includes noxious parasitic weeds from the genera *Striga, Orobanche*, and *Phelipanche* (Ishida *et al*., 2011). *Striga* species alone cause billions of dollars in annual losses by reducing yields of major crops (Pennisi, 2010). The lack of stable transgenesis protocols often hinders heritable CRISPR/Cas9 genome editing for gene function analysis in crops and species beyond standard model plants, including parasitic Orobanchaceae (Steinberger and Voytas, 2025). Here, we adapted a virus-mediated delivery system for ultracompact TnpB nucleases, enabling genome editing independently of tissue regeneration or floral dip transformation in the parasitic plant *P. japonicum* (Nagalakshmi *et al*., 2025; Weiss *et al*., 2025).

---

TnpB nucleases recognize targets through a programmable RNA guide (reRNA or ωRNA) and a transposon-associated motif (TAM) that must be present upstream of the target sequence. Expression of the nuclease and guide RNA as a single transcript fused to the hepatitis delta virus (HDV) ribozyme at the 3′ end significantly improves editing efficiency while drastically reducing size and therefore enabling virus-mediated delivery (Nagalakshmi *et al*., 2025; Weiss *et al*., 2025).

In contrast to previous virus-induced gene editing (VIGE) protocols, we used *Agrobacterium rhizogenes* to deliver the two modified Tobacco Rattle Virus (TRV) vectors into 12-day-old *P. japonicum* seedlings (Figure 1a). *Agrobacterium rhizogenes*-mediated transformation is well established for several crop and model species, including *P. japonicum* (Ishida *et al*., 2011). Using the TnpB-type genome editor Ymu1, we targeted two genes with easily scorable loss-of-function phenotypes. *Phytoene desaturase* (*PDS*) is essential for carotenoid biosynthesis and its disruption leads to photobleaching (Nagalakshmi *et al*., 2025; Weiss *et al*., 2025). *3,8-divinyl protochlorophyllide a 8-vinyl reductase* (DVR) catalyses the reduction of the 8-vinyl group of chlorophyll molecules and mutants exhibit a pale green phenotype and high chlorophyll fluorescence (Nagata *et al*., 2005). Following transformation with these constructs, *P. japonicum* developed necrotic regions, primarily in cotyledons, likely as a result of successful TRV assembly and spread (Figure S1). We used three guides targeting *PjPDS* (Pjv1_00029476) and two guides targeting *PjDVR* (Pjv1_00001967) (Figure 1b). From week two after transformation, TRV2-Ymu1-*PDS*g2-infected plants developed white sectors, indicative of *PDS* disruption (Figure 1c). Sanger sequencing confirmed editing at the *PDS* locus (Figure S2). Editing frequencies were quantified by amplicon sequencing in pooled systemic leaves that developed after transformation. While *PDS*g1 and *PDS*g3 showed little to no detectable editing, TRV2-Ymu1-*PDS*g2-infected tissue exhibited editing rates between 25–40%. Editing efficiencies for *DVRg1* and *DVRg2* were up to 10% (Figure 1f).

**FIGURE 1.**
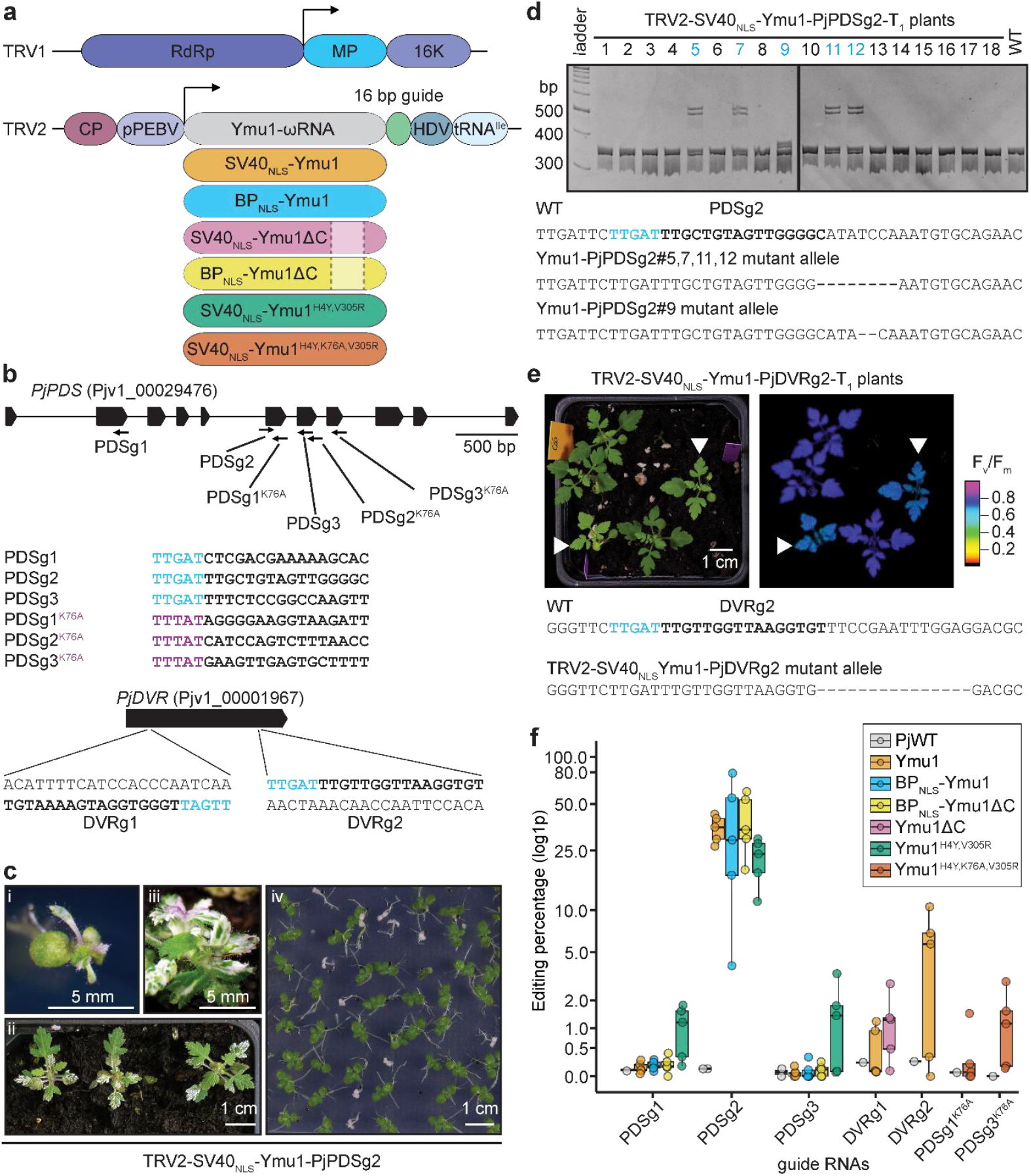
Heritable genome editing in *Phtheirospermum japonicum* using viral delivery of Ymu1. (a) Schematic of the two genomic segments of Tobacco rattle virus (TRV). Six different Ymu1 variants were inserted into the cargo region of TRV2 under the control of a pea early browning virus subgenomic promoter (pPEBV). A 16-nt guide sequence is followed by a hepatitis delta virus (HDV) ribozyme to generate homogenous 3’ ends, and a truncated tRNA^Ile^ enhances meristem entrance. (b) Genomic positions of guide RNAs targeting *PYTOENE DESATURASE* (*PDS*) and *3,8-DIVINYL PROTOCHLOROPHYLLIDE A 8-VINYL REDUCTASE* (*DVR*). The transposon-associated motif (TAM) is indicated in blue or purple for the Ymu1^K76A^ variant. (c) *PDS* knockout phenotype observed in plants infected with Ymu1 and *PjPDS*g2: (i) 18 days after transformation; (ii) after transfer to soil; (iii) seed pod with white sectors; and (iv) offspring of a heterozygous T_1_ plant. (d) Genotyping of T_1_ plants derived from a white-sectored seed pod by PCR and native polyacrylamide gel electrophoresis. Heteroduplex DNA in heterozygous plants shows reduced mobility. The vertical black line indicates where gels were merged. Mutant alleles (blue) were confirmed by Sanger sequencing and are shown below the gel. (e) Pale-green phenotype of T_1_ offspring from a Ymu1–*PjDVRg2*-infected plant (left) and corresponding chlorophyll fluorescence images (right). The mutant allele (15 bp deletion) is shown below. (f) Editing efficiencies in systemic leaves determined by amplicon sequencing for different guide RNAs and Ymu1 variants.

Several TRV2-Ymu1-*PDS*g2-infected plants with variegated sepals developed seed pods with white sectors (Figure 1c). Germinated seeds from such seed pods developed into green seedlings. In five of the 18 tested plants, native polyacrylamide gel electrophoresis showed slower-migrating PCR products resulting from heteroduplex formation of different *PDS* alleles (Figure 1d). Sanger sequencing confirmed two distinct deletion alleles at the *PDS* locus (2 bp and 8 bp, Figure 1d). Following self-fertilization, homozygous *PDS* knockout plants were recovered at a frequency of 25% in the T_2_ generation, indicating a stably inherited heterozygous *PDS* editing event in the previous generation (Figure 1c, iv). For *DVR*, biallelic double-edits were already obtained in the T_1_ generation, leading to pale-green *P. japonicum* plants with high chlorophyll fluorescence, reminiscent of *dvr* loss-of-function mutants (Figure 1e). Sanger sequencing confirmed that both alleles in these plants carried an identical 15 bp deletion, resulting in a mutant protein variant lacking five conserved amino acids (Figure S3). Collectively, these results demonstrate that virus-delivered Ymu1 can efficiently disrupt endogenous genes in a parasitic Orobanchaceae.

To improve editing efficiency, we incorporated previously described mutations into the coding sequence of *Ymu1*. Two mutations (H4Y and V305R) were reported to enhance editing activity in *E. coli* and in planta, while an additional mutation, K76A, was shown in the related TnpB enzyme Dra2 to alter TAM specificity from TTGAT to TYTAT (Marquart *et al*., 2024). We also introduced a bipartite nuclear localisation signal, which has been shown to improve genome editing efficiency of the CRISPR/Cas9 system (Develtere *et al*., 2024). Furthermore, during the cloning process, we repeatedly isolated a 103-bp deletion within a direct repeat that spans portions of the *Ymu1* coding region and the ωRNA. This deletion does not alter the amino acid sequence of Ymu1 but results in a shortened transcript (Figure S4). The same deletion is present in the original report describing Ymu1-mediated genome editing in *Arabidopsis* (Weiss *et al*., 2025). We assessed editing efficiencies of the different *Ymu1* variants by amplicon sequencing in systemic leaves. The combined H4Y and V305R mutations improved editing for the *PDS*g1 and *PDS*g3 guides, which showed no detectable activity when using the wild-type *Ymu1* sequence. In contrast, the introduction of the bipartite nuclear localisation signal had no effect on editing efficiency for any of the guides tested (Figure 1f). Interestingly, the C-terminal deletion did not impair editing activity. This shortened Ymu1–ωRNA construct may facilitate future applications by allowing the addition of functional domains while remaining within TRV2 cargo size constraints. For the *Ymu1*^K76A^ variant with altered TAM specificity, two guides (*PDS*g1^K76A^ and *PDS*g3^K76A^) showed detectable editing in at least a subset of samples (Figure 1f). Editing efficiency for *PDS*g2^K76A^ could not be reliably quantified, as wild-type PCR products also contained deletions overlapping the target site, likely due to amplification errors in repetitive sequence regions during PCR.

Given that we were able to obtain homozygous mutants for both target genes using the wild-type *Ymu1* sequence, we are confident that the improved *Ymu1* variants and the expanded TAM compatibility of the *Ymu1*^K76A^ variant will enable genome editing of many genes in *P. japonicum* and other parasitic Orobanchaceae, including candidates involved in parasitism. Beyond parasitic plants, our approach to use *Agrobacterium-rhizogenes*-mediated transformation may find broad application in crops and other (non-standard) model systems.

## Author Contributions

H.R. and T.S. conceived and designed the research. H.R. conducted the experiments and data analysis. V.Z. supported molecular cloning. H.R. and T.S. wrote the manuscript.

## Acknowledgements

The work was supported by grants of the German Research Foundation (#424122841).

## Data Availability Statement

Amplicon sequencing data were deposited at NCBI-SRA and are available under BioProject PRJNA1421118.

## Supporting Information

Additional supporting information can be found online comprising Methods S1, Figures S1– S4 and Table S1.

**Methods S1: Material and Methods**

**Figure S1: Phenotypes of TRV-infected *Phtheirospermum japonicum* plants**.

**Figure S2: Sanger sequencing of TRV2**–**Ymu1**–***PjPDS*g2-infected *Phtheirospermum japonicum* plants**.

**Figure S3: Alignment of protein sequences of 3**,**8-divinyl protochlorophyllide a 8-vinyl reductase (DVR) from six different angiosperms**.

**Figure S4: Alignment of DNA sequences for Ymu1 variants used for cloning.**

**Table S1: Oligonucleotides used in this study**

## Notes

### Competing Interest Statement

The authors have declared no competing interest.

