## Supplementary Figures for "Virus-induced genome editing in the parasitic plant *Phtheirospermum japonicum*"

### Material and Methods

#### Molecular Cloning

An internal BsaI site in TRV2 expression vector SPDK3888 (Ellison *et al.*, 2020) was first removed by amplifying the entire plasmid using Q5 polymerase (NEB). The PCR product was digested with DpnI to eliminate the methylated template plasmid, purified, and subsequently circularized using a Golden Gate assembly reaction with BsaI and T4 DNA ligase, yielding the vector SPDK3888-$BsaI_{removed}$. The coding sequence of TnpB nuclease Ymu1, fused to an N-terminal SV40 nuclear localization signal (NLS) and followed by the ωRNA and two BsaI sites for guideRNA insertion, was synthesized (Genewiz). Introduced HpaI and XmaI sites were used to clone the Ymu1- ωRNA cassette into SPDK3888-$BsaI_{removed}$. Ymu1 variants containing alternative NLS sequences or point mutations were generated by KLD mutagenesis using the synthesized plasmids as templates and subsequently cloned into SPDK3888-$BsaI_{removed}$ as described above.

For guide RNA cloning, two complementary oligonucleotides carrying 5′-TCAA (sense) or 5′-GGCC (antisense) overhangs were phosphorylated using T4 polynucleotide kinase (PNK) in T4 DNA ligase buffer in a combined reaction. The oligonucleotides were annealed by heating to 95 °C for 5 min, followed by gradual cooling to room temperature. Annealed oligonucleotides were inserted into the TRV2 vectors by Golden Gate assembly using BsaI. All oligonucleotides used in this study are listed in Table S1.

#### Plant material, growth conditions, transformation and chlorophyll fluorescence measurements

*Phtheirospermum japonicum* (Thunb.) Kanitz ecotype Okayama wild-type (WT) (Ishida *et al.*, 2011) seeds were surface-sterilized for 10 min in 70% (v/v) ethanol containing 0.05% (w/v) SDS, washed four times with 96% ethanol, and air-dried. Plants used for *Agrobacterium rhizogenes*-mediated transformation were grown on Gamborg B5 medium (Duchefa, Cat. No. G0209) supplemented with 1% (w/v) sucrose (Duchefa, Cat. No. S0809.5000) and 0.6% (w/v) agar for 12 days under a 12 h/12 h light/dark photoperiod (~100 μmol $m^{-2}$ $s^{-1}$) at 21 °C. Transformations using *Agrobacterium rhizogenes* strain AR1193 were performed according to Ishida et al. (2011), with the following modifications. *Agrobacterium* strains carrying vectors for Tobacco rattle virus (TRV) segment 1 (pYL192, Liu et al., 2002) and TRV segment 2 were mixed at a 1:1 ratio to a final $OD_{600}$ of 0.15 for each strain. Two days after transformation, plants were transferred to Gamborg B5 medium supplemented with 300 μg $mL^{-1}$ cefotaxime and grown for three weeks under a 12 h/12 h light/dark photoperiod at 21 °C before transfer to soil. $T_1$ and $T_2$ seeds were surface-sterilized as described above and germinated on Gamborg B5 medium containing 1% (w/v) sucrose. $T_1$ plants were transferred to soil after approximately three weeks, and growth was continued under a 12 h/12 h light/dark photoperiod at 21 °C. Chlorophyll fluorescence parameters ($F_v/F_m$) were measured on dark-adapted plants using an IMAGING-PAM Maxi chlorophyll fluorometer (WALZ).

#### Amplicon sequencing

Two systemic leaves per plant were collected from 2–3 plants per replicate and used for genomic DNA extraction. Leaf tissue was homogenized using a bead mill, and genomic DNA was isolated by isopropanol precipitation following lysis in extraction buffer (200 mM Tris–HCl, pH 7.5; 250 mM NaCl; 0.5% [w/v] SDS; 25 mM EDTA). After washing with ethanol, DNA was resuspended in TE buffer and used as template for PCR amplification with Q5 polymerase (NEB) and gene-specific primers listed in Table S1. PCR products were subsequently used as templates for a second PCR with barcoded primers (Table S1) to introduce inner barcodes and attachment sequences for a third PCR using outer barcodes and universal adapter sequences required for amplicon sequencing (Eurofins Genomics or Genewiz/Azenta). Up to 40 samples with unique barcode combinations were pooled, purified twice to remove residual primers, and sequenced in paired-end mode (2 × 300 bp or 2 × 250 bp).

Sequencing reads were demultiplexed using cutadapt (Martin, 2011) and analyzed with CRISPResso2 (Clement *et al.*, 2019) using the parameters --ignore_substitutions and a quantification window of 10 bp.

**Native PAGE analysis of PCR products and Sanger sequencing**

Genomic DNA was isolated from $T_1$ plants as described for amplicon sequencing and used as template for PCR amplification with Q5 polymerase (NEB) and primers listed in Table S1. PCR products were resolved on 8% native polyacrylamide gels prepared in TBE buffer. Gels were stained with HDGreen Plus (INTAS) and visualized under UV illumination. PCR products were subsequently purified and submitted for Sanger sequencing using the reverse primer.

#### Supplementary Figures

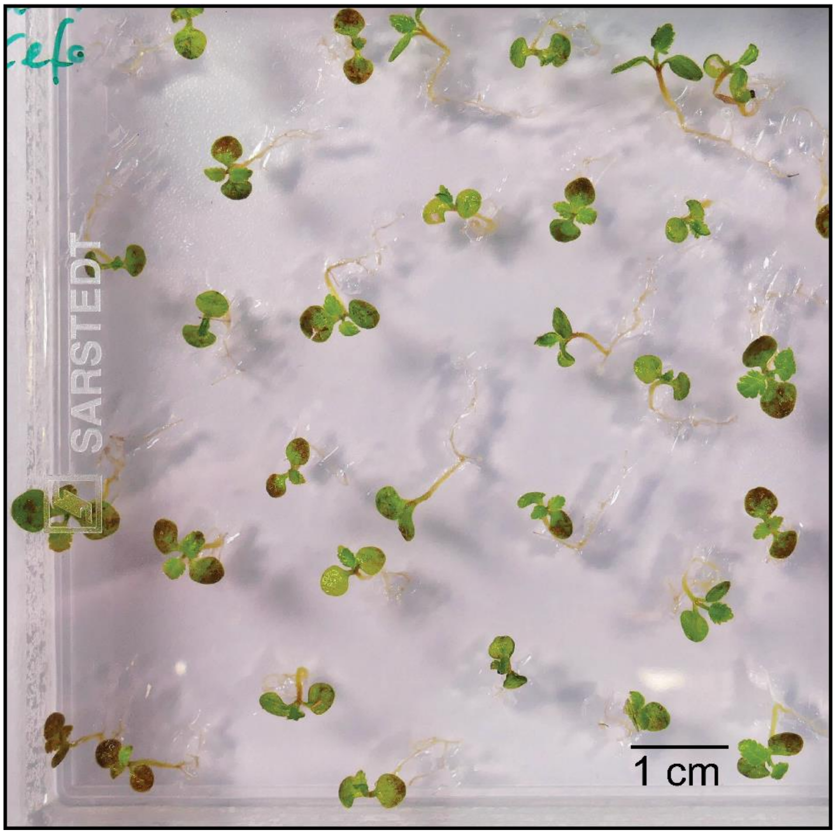


**Figure S1 | Phenotypes of TRV-infected *Phtheirospermum japonicum* plants.** Phenotype of *Phtheirospermum japonicum* plants 8 days after transformations with two *Agrobacterium rhizogenes* AR1193 strains expressing TRV1 and TRV2-Ymu1-*PjDVR*g1. Many cotyledons show brown discoloration indicative of virus-induced necrosis.

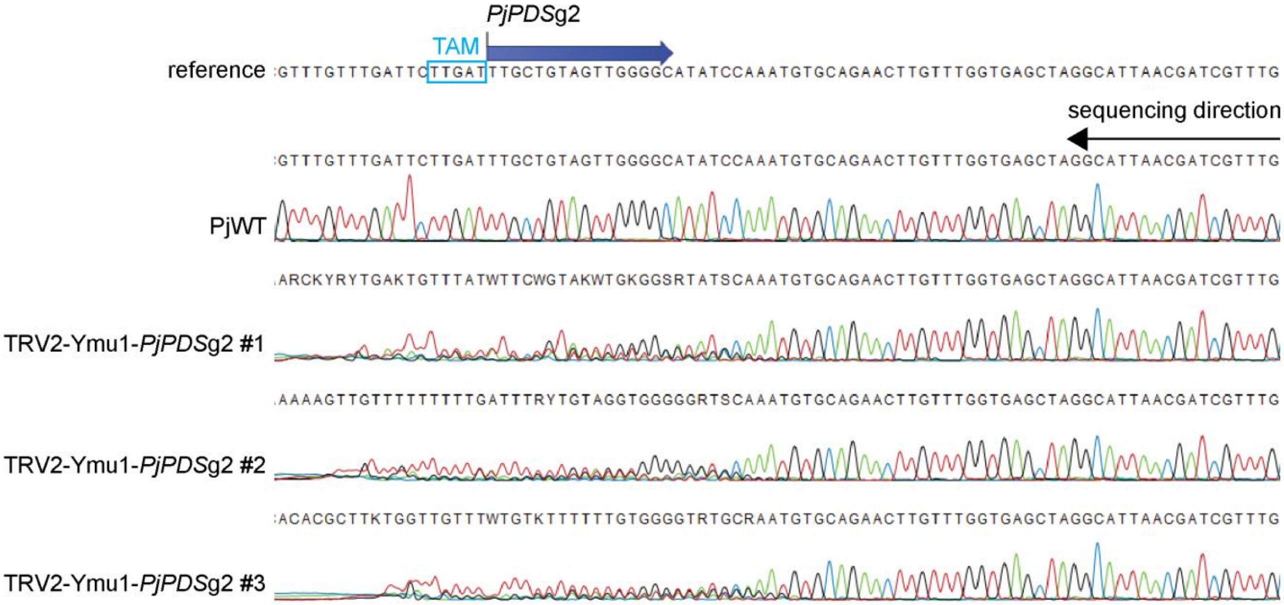


**Figure S2 | Sanger sequencing of TRV2–Ymu1–PjPDSg2-infected *Phtheirospermum japonicum* plants**. A systemic leaf was harvested from each of three plants 17 days after inoculation with TRV1 and TRV2–Ymu1–*PjPDS*g2. Genomic DNA was isolated, and the region surrounding the target site was amplified by PCR. PCR products were purified and sequenced using the reverse primer. TRV2–Ymu1–*PjPDS*g2-infected plants display polymorphic sequences at the guide RNA target site. The 16-nt guide RNA sequence is indicated by a dark blue arrow above the reference sequence. The transposon-associated motif (TAM) required upstream of the guide RNA target site is highlighted by a light blue box.

```
ZmDVR      MATILLSS------RLPT------------TGTATPSPTRPAPRFLSFP----GTAIRRR     38
AtDVR      MSLCSSFNVFASYSPKPK-------TIFKDSKFISQFQVKSSPLASTFHTNESSTSLKYK     53
PjDVR      MSICTSSFNCN--GLTLHSSKAHTFKRVFTSQFITKSQVTTIPYAIPFPSLYFSKPFKIS     58
VvDVR      MSLYLSSNVLTLHSP-----KTRSFRNCSASQFINQNQVTPAPYAITRLPLSLSQSPKFS     55
MtDVR      MSLCYSSSTFITHPSLKHQTLSFNFSSHPPSHFINLFKVK------------SNRPIKYT     48
PvDVR      MSLCYTSNFIS-LNH--QKSLSLTFSSDS-PRFINLFSVKHRK---------PHHPIKFT     47
           *:                                .   .                  :

ZmDVR      GRGPLLASSAVSPPAPASAAQPYRALPASETTVLVTGATGYIGRYVVWELLRRGHRVLAV     98
AtDVR      RARLKPISSLDSGISEIATSPSFRNKSPKDINVLVVGSTGYIGRFVVKEMIKRGFNVIAV     113
PjDVR      KKRLNLVT-FSSTQSIETPKPSFRTKNPKDINVLVVGSTGYIGKFVVKELIKRGFNVIAI     117
VvDVR      RERFL--PITASITPTVEPPSSFRGKNASEINVVVVGSTGYIGKFVVKELVSRGFNVIAI     113
MtDVR      KQKLKLYASLSQSEQIETTPTTFRIKNPKDVNVLVVGSTGYIGKFVVKELIQRGFNVTAI     108
PvDVR      TERFKLVASLTPAPPIETAPSSYRSKSPKDVNVLVVGSTGYIGKYVVRELVKRGFNVTAI     107
                                 :*    .: .*:*.*:*****::** *:: **..* *:

ZmDVR      ARSRSGIRGRNSPDDVVADLAPAQVVFSDVTDPAALLADLAPHG-PVHAAVCCLASRGGG     157
AtDVR      AREKSGIRGKNDKEETLKQLQGANVCFSDVTELDVLEKSIENLGFGVDVVVSCLASRNGG     173
PjDVR      AREKSGIKGKNSKDETLGMLTGANVCFSDVTDLSSLSKTIEGLETPIDVVVSCLASRSGG     177
VvDVR      ARERSGIRGRNRKEDTLTELNGANVWFSDVTSLDVLEKSLENLGLPIDVVVSCLASRTGG     173
MtDVR      AREKSGIKGSIDKETTLNELRGANVCFSDVTNLDVFDEDLKNLGVGFDVVVSCLASRNGG     168
PvDVR      ARERSGIKGSVDKDQTLNQLRGANVCFSDVANLDALEGSLNSLGNSFDVVVSCLASRNGG     167
           **.:***:*    : .:  *  *:* ****:.   :   :      ....*.***** **

ZmDVR      VQDSWRVDYRATLHTLQAARGLGAAHFVLLSAICVQKPLLEFQRAKLKFEEELAAEAARD     217
AtDVR      IKDSWKIDYEATKNSLVAGKKFGAKHFVLLSAICVQKPLLEFQRAKLKFEAELMDLAEQQ     233
PjDVR      VKDSWLIDYQATKNSLVAGRKFGAAHFVLLSAICVQKPLLEFQRAKLKFEAELIQEAENG     237
VvDVR      VKDSWKIDYEATKNSLVAGRKRGASHFVLLSAVCVQKPLLEFQRAKLKFEAELMKEAEED     233
MtDVR      VKDSWKIDYEATKNSLLAGRKLGASHFVLLSAICVQKPLLEFQRAKLKLEDELVKEAEKD     228
PvDVR      VKDSWKIDYEATRNSLVAGRKRGASHFVLLSAICVQKPLLEFQRAKLKFEDELMKLAEED     227
           ::*** :**.** ::* *.:  ** *******:**************:* **   * .

ZmDVR      P-SFTYSVVRPTAFFKSLGGQVDIVKNGQPYVMFGDGKLCACKPISEEDLAAFIADCIYD     276
AtDVR      DSSFTYSIVRPTAFFKSLGGQVEIVKDGKPYVMFGDGKLCACKPISEQDLAAFIADCVLE     293
PjDVR      NGLFTYSIVRPTAFFKSLAGQVELVKDGKPYVMFGDGKLCACKPISEADLASFIADCLLS     297
VvDVR      D-GFTYSIVRPTAFFKSLGGQVELVKDGKPYVMFGDGKLCACKPISEQDLASFIADCVLE     292
MtDVR      D-RFSYSIVRPTAFFKSLGGQVDLVKDGKPYVMFGDGKLCACKPISEQDLASFIVDCVMS     287
PvDVR      G-GFSYSIVRPTAFFKSLGGQVELVKDGKPYVMFGDGKLCACKPISESDLASFIVDCVLS     286
              *:**:*********.***::**:*:***************** ***:**.**: .

ZmDVR      QDKANKVLPIGGPGKALTPLEQGEMLFRLLGREPKFIKVPIQIMDAVIWVLDGLAKLFPG     336
AtDVR      ENKINQVLPIGGPGKALTPLEQGEILFKILGREPKFLKVPIEIMDFVIGVLDSIAKIFPS     353
PjDVR      EGLVNRILPVGGPGEALTPLEQGEMLFRLVEREPKFLKVPIQIMDFAIGVLDLLVKVFPN     357
VvDVR      KDKINQVLPIGGPGKALTPLEQGEMLFRLAGRKPNFLKVPIGIMDFAIGFLDFLVKIFPS     352
MtDVR      EDKINKILPIGGPGKALTPLEQGEILFKLLRKEPKFLKVPIGIMDFAIGVLDNLVKVFPS     347
PvDVR      EDKINKVLPIGGPGKALTPLEQGEILFRLLGKEPKFLKVPIGIMDFAIGVLDFLVKVFPS     346
           :.  *::**:****:*********:**::  ::*:*:**** *** .* .** :.*:**.

ZmDVR      LEDAAEFGKIGRYYASESMLLLDPETGEYSDEKTPSYGKDTLEQFFQRVIREGMAGQELG     396
AtDVR      VGEAAEFGKIGRYYAAESMLILDPETGEYSEEKTPSYGKDTLEDFFAKVIREGMAGQELG     413
PjDVR      LEDAAEFGKIGRYYAAESMLVWDEERGEYDAESTPSYGEDTLEEFFRRVLEEGMAGQELG     417
VvDVR      MEDAAEFGKIGRYYAAESMLVLDPETGEYSAEKTPSYGKDTLEEFFERVLREGMAGQELG     412
MtDVR      LEDAAEFGKIGRYYAAESMLILDPDTGEYSDEKTPSYGNDTLEDFFARVLREGMAGQELG     407
PvDVR      LEDAAEFGKIGRYYAAESMLLLDPETGEYSAEKTPSYGNDTLEEFFATVLREGMAGQELG     406
           : :***********:****: * : ***. *.*****:****:**  *:.*********

ZmDVR      EQTI--  400
AtDVR      EQFF--  417
PjDVR      EQAIF*  422
VvDVR      EQTIF-  417
MtDVR      EQTIF-  412
PvDVR      EQTIF-  411
           ** :
```

**Figure S3 | Alignment of protein sequences of 3,8-divinyl protochlorophyllide a 8-vinyl reductase (DVR) from six different angiosperms.** Protein sequences were obtained by Protein BLAST using the *Arabidopsis thaliana* DVR protein as the query. The five amino acids deleted in TRV2–Ymu1–PjDVRg2 plants are highlighted in magenta. Abbreviations: Zm, *Zea mays*; At, *Arabidopsis thaliana*; Pj, *Phtheirospermum japonicum*; Vv, *Vitis vinifera*; Mt, *Medicago truncatula*; Pv, *Phaseolus vulgaris*.

```
HpaI recognition sequence     Start
SV40NLS-Ymu1-WT            GTTAACGAGATGGCTTCTTCTCCT------------------------------CCTAAG
SV40NLS-Ymu1-H4Y,V305R     GTTAACGAGATGGCTTCTTCTCCT------------------------------CCTAAG
BPNLS-Ymu1                 GTTAACGAGATGGCTTCTTCTCCTAAGAGAACTGCTGATGGATCTGAGTTCGAACCTAAG
SV40NLS-Ymu1-deltaC        GTTAACGAGATGGCTTCTTCTCCT------------------------------CCTAAG
Ymu1-H4Y,K76A,V305R        GTTAACGAGATGGCTTCTTCTCCT------------------------------CCTAAG

                           ************************                              ******

SV40NLS-Ymu1-WT            AAAAAGAGAAAGGTTTCTTGGAAGCTGCAGCATAAAGCCTATGAATACCGTATCTATCCA
SV40NLS-Ymu1-H4Y,V305R     AAAAAGAGAAAGGTTTCTTGGAAGCTGCAGTATAAAGCCTATGAATACCGTATCTATCCA
BPNLS-Ymu1                 AAAAAGAGAAAGGTTTCTTGGAAGCTGCAGCATAAAGCCTATGAATACCGTATCTATCCA
SV40NLS-Ymu1-deltaC        AAAAAGAGAAAGGTTTCTTGGAAGCTGCAGCATAAAGCCTATGAATACCGTATCTATCCA
Ymu1-H4Y,K76A,V305R        AAAAAGAGAAAGGTTTCTTGGAAGCTGCAGTATAAAGCCTATGAATACCGTATCTATCCA
                           ****************************** *****************************

SV40NLS-Ymu1-WT            GATAAGAAGCAGGAAACACTGATTGCCAAGACAATAGGCAGTTCAAGGTATGTATACAAC
SV40NLS-Ymu1-H4Y,V305R     GATAAGAAGCAGGAAACACTGATTGCCAAGACAATAGGCAGTTCAAGGTATGTATACAAC
BPNLS-Ymu1                 GATAAGAAGCAGGAAACACTGATTGCCAAGACAATAGGCAGTTCAAGGTATGTATACAAC
SV40NLS-Ymu1-deltaC        GATAAGAAGCAGGAAACACTGATTGCCAAGACAATAGGCAGTTCAAGGTATGTATACAAC
Ymu1-H4Y,K76A,V305R        GATAAGAAGCAGGAAACACTGATTGCCAAGACAATAGGCAGTTCAAGGTATGTATACAAC
                           ************************************************************

SV40NLS-Ymu1-WT            CATTTCCTGGAACTTTGGAACAAGGAATATGAGGAGACTGGAAAAGGTCTGACTTATTAC
SV40NLS-Ymu1-H4Y,V305R     CATTTCCTGGAACTTTGGAACAAGGAATATGAGGAGACTGGAAAAGGTCTGACTTATTAC
BPNLS-Ymu1                 CATTTCCTGGAACTTTGGAACAAGGAATATGAGGAGACTGGAAAAGGTCTGACTTATTAC
SV40NLS-Ymu1-deltaC        CATTTCCTGGAACTTTGGAACAAGGAATATGAGGAGACTGGAAAAGGTCTGACTTATTAC
Ymu1-H4Y,K76A,V305R        CATTTCCTGGAACTTTGGAACAAGGAATATGAGGAGACTGGAAAAGGTCTGACTTATTAC
                           ************************************************************

SV40NLS-Ymu1-WT            GCATGCTCCAAACTTCTGACAAAGCTTAAGAGGGACCCGGAAACAGTATGGCTTTGTGAA
SV40NLS-Ymu1-H4Y,V305R     GCATGCTCCAAACTTCTGACAAAGCTTAAGAGGGACCCGGAAACAGTATGGCTTTGTGAA
BPNLS-Ymu1                 GCATGCTCCAAACTTCTGACAAAGCTTAAGAGGGACCCGGAAACAGTATGGCTTTGTGAA
SV40NLS-Ymu1-deltaC        GCATGCTCCAAACTTCTGACAAAGCTTAAGAGGGACCCGGAAACAGTATGGCTTTGTGAA
Ymu1-H4Y,K76A,V305R        GCATGCTCCAAACTTCTGACAAAGCTTAAGAGGGACCCGGAAACAGTATGGCTTTGTGAA
                           ************************************************************

SV40NLS-Ymu1-WT            GTGGATAAGTTCTCGCTCCAGAACTCGCTTCGTAACCTGTCGGACGCCTTCTCCCGCTTT
SV40NLS-Ymu1-H4Y,V305R     GTGGATAAGTTCTCGCTCCAGAACTCGCTTCGTAACCTGTCGGACGCCTTCTCCCGCTTT
BPNLS-Ymu1                 GTGGATAAGTTCTCGCTCCAGAACTCGCTTCGTAACCTGTCGGACGCCTTCTCCCGCTTT
SV40NLS-Ymu1-deltaC        GTGGATAAGTTCTCGCTCCAGAACTCGCTTCGTAACCTGTCGGACGCCTTCTCCCGCTTT
Ymu1-H4Y,K76A,V305R        GTGGATGCGTTCTCGCTCCAGAACTCGCTTCGTAACCTGTCGGACGCCTTCTCCCGCTTT
                           ******  ****************************************************

SV40NLS-Ymu1-WT            TTCAAAGGACAGAACGAGCATCCCCAGTTCAAGAGCAAGAAGAGCCCGAGGCAAAGCTAC
SV40NLS-Ymu1-H4Y,V305R     TTCAAAGGACAGAACGAGCATCCCCAGTTCAAGAGCAAGAAGAGCCCGAGGCAAAGCTAC
BPNLS-Ymu1                 TTCAAAGGACAGAACGAGCATCCCCAGTTCAAGAGCAAGAAGAGCCCGAGGCAAAGCTAC
SV40NLS-Ymu1-deltaC        TTCAAAGGACAGAACGAGCATCCCCAGTTCAAGAGCAAGAAGAGCCCGAGGCAAAGCTAC
Ymu1-H4Y,K76A,V305R        TTCAAAGGACAGAACGAGCATCCCCAGTTCAAGAGCAAGAAGAGCCCGAGGCAAAGCTAC
                           ************************************************************

SV40NLS-Ymu1-WT            ACGACACAATATACGAACAACAACATCGCAGTTTCCGGAAATTGTCTGAAGCTGCCTAAG
SV40NLS-Ymu1-H4Y,V305R     ACGACACAATATACGAACAACAACATCGCAGTTTCCGGAAATTGTCTGAAGCTGCCTAAG
BPNLS-Ymu1                 ACGACACAATATACGAACAACAACATCGCAGTTTCCGGAAATTGTCTGAAGCTGCCTAAG
SV40NLS-Ymu1-deltaC        ACGACACAATATACGAACAACAACATCGCAGTTTCCGGAAATTGTCTGAAGCTGCCTAAG
Ymu1-H4Y,K76A,V305R        ACGACACAATATACGAACAACAACATCGCAGTTTCCGGAAATTGTCTGAAGCTGCCTAAG
                           ************************************************************

SV40NLS-Ymu1-WT            CTGGGCCTTGTCAAGTTTGCAGACAGTAGGGAGATGAAAGGCCGTATACTGAATGCCACA
SV40NLS-Ymu1-H4Y,V305R     CTGGGCCTTGTCAAGTTTGCAGACAGTAGGGAGATGAAAGGCCGTATACTGAATGCCACA
BPNLS-Ymu1                 CTGGGCCTTGTCAAGTTTGCAGACAGTAGGGAGATGAAAGGCCGTATACTGAATGCCACA
SV40NLS-Ymu1-deltaC        CTGGGCCTTGTCAAGTTTGCAGACAGTAGGGAGATGAAAGGCCGTATACTGAATGCCACA
Ymu1-H4Y,K76A,V305R        CTGGGCCTTGTCAAGTTTGCAGACAGTAGGGAGATGAAAGGCCGTATACTGAATGCCACA
                           ************************************************************
```

```
SV40NLS-Ymu1-WT        GTACGAAGGAAATCGAGCGGGAAATTCTTCGTATCGATCCTATGCGAAGAGGAGATCTGT
SV40NLS-Ymu1-H4Y,V305R GTACGAAGGAAATCGAGCGGGAAATTCTTCGTATCGATCCTATGCGAAGAGGAGATCTGT
BPNLS-Ymu1             GTACGAAGGAAATCGAGCGGGAAATTCTTCGTATCGATCCTATGCGAAGAGGAGATCTGT
SV40NLS-Ymu1-deltaC    GTACGAAGGAAATCGAGCGGGAAATTCTTCGTATCGATCCTATGCGAAGAGGAGATCTGT
Ymu1-H4Y,K76A,V305R    GTACGAAGGAAATCGAGCGGGAAATTCTTCGTATCGATCCTATGCGAAGAGGAGATCTGT
                       ************************************************************

SV40NLS-Ymu1-WT        GAATTGCCAAAGACTGACTCATCTGTCGGAATTGACCTCGGAATCATTGACTTTGCGGTT
SV40NLS-Ymu1-H4Y,V305R GAATTGCCAAAGACTGACTCATCTGTCGGAATTGACCTCGGAATCATTGACTTTGCGGTT
BPNLS-Ymu1             GAATTGCCAAAGACTGACTCATCTGTCGGAATTGACCTCGGAATCATTGACTTTGCGGTT
SV40NLS-Ymu1-deltaC    GAATTGCCAAAGACTGACTCATCTGTCGGAATTGACCTCGGAATCATTGACTTTGCGGTT
Ymu1-H4Y,K76A,V305R    GAATTGCCAAAGACTGACTCATCTGTCGGAATTGACCTCGGAATCATTGACTTTGCGGTT
                       ************************************************************

SV40NLS-Ymu1-WT        ATGTCAGACGGAAGCAGGCATGACAACAATCATTTTACCAGGCAAATGGAAGAAAGGCTT
SV40NLS-Ymu1-H4Y,V305R ATGTCAGACGGAAGCAGGCATGACAACAATCATTTTACCAGGCAAATGGAAGAAAGGCTT
BPNLS-Ymu1             ATGTCAGACGGAAGCAGGCATGACAACAATCATTTTACCAGGCAAATGGAAGAAAGGCTT
SV40NLS-Ymu1-deltaC    ATGTCAGACGGAAGCAGGCATGACAACAATCATTTTACCAGGCAAATGGAAGAAAGGCTT
Ymu1-H4Y,K76A,V305R    ATGTCAGACGGAAGCAGGCATGACAACAATCATTTTACCAGGCAAATGGAAGAAAGGCTT
                       ************************************************************

SV40NLS-Ymu1-WT        AGACGGGAGCAGCGAAAGCTTGCAAGGCGTGCACTTGCTGCGGAGAAAAGGGGCATTTCC
SV40NLS-Ymu1-H4Y,V305R AGACGGGAGCAGCGAAAGCTTGCAAGGCGTGCACTTGCTGCGGAGAAAAGGGGCATTTCC
BPNLS-Ymu1             AGACGGGAGCAGCGAAAGCTTGCAAGGCGTGCACTTGCTGCGGAGAAAAGGGGCATTTCC
SV40NLS-Ymu1-deltaC    AGACGGGAGCAGCGAAAGCTTGCAAGGCGTGCACTTGCTGCGGAGAAAAGGGGCATTTCC
Ymu1-H4Y,K76A,V305R    AGACGGGAGCAGCGAAAGCTTGCAAGGCGTGCACTTGCTGCGGAGAAAAGGGGCATTTCC
                       ************************************************************

SV40NLS-Ymu1-WT        CTTTCTGAAGCCAGGAACTATCAGAAGCAAAGGCGGAAGGTAGCGAGACTTTATGAAAAG
SV40NLS-Ymu1-H4Y,V305R CTTTCTGAAGCCAGGAACTATCAGAAGCAAAGGCGGAAGGTAGCGAGACTTTATGAAAAG
BPNLS-Ymu1             CTTTCTGAAGCCAGGAACTATCAGAAGCAAAGGCGGAAGGTAGCGAGACTTTATGAAAAG
SV40NLS-Ymu1-deltaC    CTTTCTGAAGCCAGGAACTATCAGAAGCAAAGGCGGAAGGTAGCGAGACTTTATGAAAAG
Ymu1-H4Y,K76A,V305R    CTTTCTGAAGCCAGGAACTATCAGAAGCAAAGGCGGAAGGTAGCGAGACTTTATGAAAAG
                       ************************************************************

SV40NLS-Ymu1-WT        GTTGCAAACCAGCGAAAAGAATATCTTAACAAACTCAGTACAGAGATAGTCAAAAACCAC
SV40NLS-Ymu1-H4Y,V305R GTTGCAAACCAGCGAAAAGAATATCTTAACAAACTCAGTACAGAGATAGTCAAAAACCAC
BPNLS-Ymu1             GTTGCAAACCAGCGAAAAGAATATCTTAACAAACTCAGTACAGAGATAGTCAAAAACCAC
SV40NLS-Ymu1-deltaC    GTTGCAAACCAGCGAAAAGAATATCTTAACAAACTCAGTACAGAGATAGTCAAAAACCAC
Ymu1-H4Y,K76A,V305R    GTTGCAAACCAGCGAAAAGAATATCTTAACAAACTCAGTACAGAGATAGTCAAAAACCAC
                       ************************************************************

SV40NLS-Ymu1-WT        GATATCATCTGTATCGAGGACCTTAACGTTAAGGGCATGATGCGCAATCATAAACTGGCA
SV40NLS-Ymu1-H4Y,V305R GATATCATCTGTATCGAGGACCTTAACGTTAAGGGCATGATGCGCAATCATAAACTGGCA
BPNLS-Ymu1             GATATCATCTGTATCGAGGACCTTAACGTTAAGGGCATGATGCGCAATCATAAACTGGCA
SV40NLS-Ymu1-deltaC    GATATCATCTGTATCGAGGACCTTAACGTTAAGGGCATGATGCGCAATCATAAACTGGCA
Ymu1-H4Y,K76A,V305R    GATATCATCTGTATCGAGGACCTTAACGTTAAGGGCATGATGCGCAATCATAAACTGGCA
                       ************************************************************

SV40NLS-Ymu1-WT        AAAAGCATCTCTGATGTATCATGGACGAGCCTTGTATCGAAACTGCAGTACAAGGCTTCC
SV40NLS-Ymu1-H4Y,V305R AAAAGCATCTCTGATGTATCATGGACGAGCCTTCGATCGAAACTGCAGTACAAGGCTTCC
BPNLS-Ymu1             AAAAGCATCTCTGATGTATCATGGACGAGCCTTGTATCGAAACTGCAGTACAAGGCTTCC
SV40NLS-Ymu1-deltaC    AAAAGCATCTCTGATGTATCATGGACGAGCCTTGTATCGAAACTGCAGTACAAGGCTTCC
Ymu1-H4Y,K76A,V305R    AAAAGCATCTCTGATGTATCATGGACGAGCCTTCGATCGAAACTGCAGTACAAGGCTTCC
                       *********************************  *************************

SV40NLS-Ymu1-WT        TGGTATGGAAAAGAAGTCATCAGGATAAGCAGATGGTTTCCGTCAAGTCAGATATGCTCA
SV40NLS-Ymu1-H4Y,V305R TGGTATGGAAAAGAAGTCATCAGGATAAGCAGATGGTTTCCGTCAAGTCAGATATGCTCA
BPNLS-Ymu1             TGGTATGGAAAAGAAGTCATCAGGATAAGCAGATGGTTTCCGTCAAGTCAGATATGCTCA
SV40NLS-Ymu1-deltaC    TGGTATGGAAAAGAAGTCATCAGGATAAGCAGATGGTTTCCGTCAAGTCAGATATGCTCA
Ymu1-H4Y,K76A,V305R    TGGTATGGAAAAGAAGTCATCAGGATAAGCAGATGGTTTCCGTCAAGTCAGATATGCTCA
                       ************************************************************
```

```
SV40NLS-Ymu1-WT         GAGTGCGGCCACAAGGACAGGAAGAAGCCTCTCCATGTAAGGGAGTGGACCTGTCCTGTT
SV40NLS-Ymu1-H4Y,V305R  GAGTGCGGCCACAAGGACAGGAAGAAGCCTCTCCATGTAAGGGAGTGGACCTGTCCTGTT
BPNLS-Ymu1              GAGTGCGGCCACAAGGACAGGAAGAAGCCTCTCCATGTAAGGGAGTGGACCTGTCCTGTT
SV40NLS-Ymu1-deltaC     GAGTGCGGCCACAAGGACAGGAAGAAGCCTCTCCATGTAAGGGAGTGGACCTGTCCTGTT
Ymu1-H4Y,K76A,V305R     GAGTGCGGCCACAAGGACAGGAAGAAGCCTCTCCATGTAAGGGAGTGGACCTGTCCTGTT
                        ************************************************************

SV40NLS-Ymu1-WT         TGTCATGCCCACCATGACCGTGACGTCAATGCAGCAAGAAATATACTGGCTGAGGGACTC
SV40NLS-Ymu1-H4Y,V305R  TGTCATGCCCACCATGACCGTGACGTCAATGCAGCAAGAAATATACTGGCTGAGGGACTC
BPNLS-Ymu1              TGTCATGCCCACCATGACCGTGACGTCAATGCAGCAAGAAATATACTGGCTGAGGGACTC
SV40NLS-Ymu1-deltaC     TGTCATGCCCACCATGACCGTGACGTCAATGCAGCAAGAAATATACTGGCTGAGGGACTC
Ymu1-H4Y,K76A,V305R     TGTCATGCCCACCATGACCGTGACGTCAATGCAGCAAGAAATATACTGGCTGAGGGACTC
                        ************************************************************
                                                   Stop
SV40NLS-Ymu1-WT         AGGATAAGAGCTCTGACTCCGGGGTCTTGAGAGATCTTTGACAGCTAGCTCAGTCCTAGG
SV40NLS-Ymu1-H4Y,V305R  AGGATAAGAGCTCTGACTCCGGGGTCTTGAGAGATCTTTGACAGCTAGCTCAGTCCTAGG
BPNLS-Ymu1              AGGATAAGAGCTCTGACTCCGGGGTCTTGAGAGATCTTTGACAGCTAGCTCAGTCCTAGG
SV40NLS-Ymu1-deltaC     AGGATAAGAGCTCTGACTCCGGGGTCTT--------------------------------
Ymu1-H4Y,K76A,V305R     AGGATAAGAGCTCTGACTCCGGGGTCTTGAGAGATCTTTGACAGCTAGCTCAGTCCTAGG
                        ****************************

SV40NLS-Ymu1-WT         TATAATAGTCAATGCAGCAAGAAATATACTGGCTGAGGGACTCAGGATAAGAGCTCTGAC
SV40NLS-Ymu1-H4Y,V305R  TATAATAGTCAATGCAGCAAGAAATATACTGGCTGAGGGACTCAGGATAAGAGCTCTGAC
BPNLS-Ymu1              TATAATAGTCAATGCAGCAAGAAATATACTGGCTGAGGGACTCAGGATAAGAGCTCTGAC
SV40NLS-Ymu1-deltaC     ------------------------------------------------------------
Ymu1-H4Y,K76A,V305R     TATAATAGTCAATGCAGCAAGAAATATACTGGCTGAGGGACTCAGGATAAGAGCTCTGAC


SV40NLS-Ymu1-WT         TCCGGGGTCTTAGCAATACAGAAGCAAACAGGAACCGCAGGAATTGCGGGGGTAGCTTGG
SV40NLS-Ymu1-H4Y,V305R  TCCGGGGTCTTAGCAATACAGAAGCAAACAGGAACCGCAGGAATTGCGGGGGTAGCTTGG
BPNLS-Ymu1              TCCGGGGTCTTAGCAATACAGAAGCAAACAGGAACCGCAGGAATTGCGGGGGTAGCTTGG
SV40NLS-Ymu1-deltaC     -----------AGCAATACAGAAGCAAACAGGAACCGCAGGAATTGCGGGGGTAGCTTGG
Ymu1-H4Y,K76A,V305R     TCCGGGGTCTTAGCAATACAGAAGCAAACAGGAACCGCAGGAATTGCGGGGGTAGCTTGG
                                   *************************************************

SV40NLS-Ymu1-WT         TAAACAAGAGAAACCTCTGCCGGCAAAGAAATAAGCCGGTAAGTATGCTCTGTTCCCAAG
SV40NLS-Ymu1-H4Y,V305R  TAAACAAGAGAAACCTCTGCCGGCAAAGAAATAAGCCGGTAAGTATGCTCTGTTCCCAAG
BPNLS-Ymu1              TAAACAAGAGAAACCTCTGCCGGCAAAGAAATAAGCCGGTAAGTATGCTCTGTTCCCAAG
SV40NLS-Ymu1-deltaC     TAAACAAGAGAAACCTCTGCCGGCAAAGAAATAAGCCGGTAAGTATGCTCTGTTCCCAAG
Ymu1-H4Y,K76A,V305R     TAAACAAGAGAAACCTCTGCCGGCAAAGAAATAAGCCGGTAAGTATGCTCTGTTCCCAAG+-
                        ************************************************************

SV40NLS-Ymu1-WT         AATCTCGTGACTTTAGTCATGAGAGTTTCAACGAGACCAAGGATCCAAGGTCTCAGGCCG
SV40NLS-Ymu1-H4Y,V305R  AATCTCGTGACTTTAGTCATGAGAGTTTCAACGAGACCAAGGATCCAAGGTCTCAGGCCG
BPNLS-Ymu1              AATCTCGTGACTTTAGTCATGAGAGTTTCAACGAGACCAAGGATCCAAGGTCTCAGGCCG
SV40NLS-Ymu1-deltaC     AATCTCGTGACTTTAGTCATGAGAGTTTCAACGAGACCAAGGATCCAAGGTCTCAGGCCG
Ymu1-H4Y,K76A,V305R     AATCTCGTGACTTTAGTCATGAGAGTTTCAACGAGACCAAGGATCCAAGGTCTCAGGCCG
                        ************************************************************

SV40NLS-Ymu1-WT         GCATGGTCCCAGCCTCCTCGCTGGCGCCGGCTGGGCAACATGCTTCGGCATGGCGAATGG
SV40NLS-Ymu1-H4Y,V305R  GCATGGTCCCAGCCTCCTCGCTGGCGCCGGCTGGGCAACATGCTTCGGCATGGCGAATGG
BPNLS-Ymu1              GCATGGTCCCAGCCTCCTCGCTGGCGCCGGCTGGGCAACATGCTTCGGCATGGCGAATGG
SV40NLS-Ymu1-deltaC     GCATGGTCCCAGCCTCCTCGCTGGCGCCGGCTGGGCAACATGCTTCGGCATGGCGAATGG
Ymu1-H4Y,K76A,V305R     GCATGGTCCCAGCCTCCTCGCTGGCGCCGGCTGGGCAACATGCTTCGGCATGGCGAATGG
                        ************************************************************

SV40NLS-Ymu1-WT         GACCCGTAGCTCAGTTGGTTAGAGCGTTGGTCTTATGAGCCGAAGGTCGCGGGTTCGAGC
SV40NLS-Ymu1-H4Y,V305R  GACCCGTAGCTCAGTTGGTTAGAGCGTTGGTCTTATGAGCCGAAGGTCGCGGGTTCGAGC
BPNLS-Ymu1              GACCCGTAGCTCAGTTGGTTAGAGCGTTGGTCTTATGAGCCGAAGGTCGCGGGTTCGAGC
SV40NLS-Ymu1-deltaC     GACCCGTAGCTCAGTTGGTTAGAGCGTTGGTCTTATGAGCCGAAGGTCGCGGGTTCGAGC
Ymu1-H4Y,K76A,V305R     GACCCGTAGCTCAGTTGGTTAGAGCGTTGGTCTTATGAGCCGAAGGTCGCGGGTTCGAGC
                        ************************************************************
```

```
SV40NLS-Ymu1-WT        CCCGCCGGAAGCACCCGGG
SV40NLS-Ymu1-H4Y,V305R CCCGCCGGAAGCACCCGGG
BPNLS-Ymu1             CCCGCCGGAAGCACCCGGG
SV40NLS-Ymu1-deltaC    CCCGCCGGAAGCACCCGGG
Ymu1-H4Y,K76A,V305R    CCCGCCGGAAGCACCCGGG
                       *******************
```

XmaI recognition sequence

**Figure S4 | Alignment of DNA sequences of Ymu1 variants used for cloning.** The alignment shows DNA sequences used for cloning Ymu1 variants into the TRV2 expression vector SPDK3888-BsaI$_{remove}$. HpaI and XmaI recognition sites at the 5′ and 3′ ends of the sequences are highlighted in yellow. The start codon is underlined, and the stop codon is highlighted in red. The insertion that generates a more efficient bipartite nuclear localization sequence is shown in magenta. Mutations leading to amino acid changes are highlighted in green in both the sequence and the corresponding variant name, next to the mutation. A 103 bp deletion present in the Ymu1ΔC variant is highlighted in gray. BsaI recognition sites are highlighted in light blue, and the single-stranded overhangs adjacent to the BsaI sites, which hybridize with the single-stranded overlaps of the annealed oligonucleotides for guide insertion, are underlined.

#### Supplementary Table

**Table S1: Oligonucleotides used in this study**

| Primer name | Sequence | purpose |
|---|---|---|
| pDK3888_BsaIremove_for | TTGGTCTCAGCTTCTTTCCTGTGGATAGCACGTAC | BsaI site removal |
| pDK3888_BsaIremove_rev | TTGGTCTCGAAGCTTTTTCGACCTTTTTCCCCTGCTA | BsaI site removal |
| Ymu1_BPinsertion_for | ggatctgagttcgaaCCTAAGAAAAAGAGAAAGGTTTC | KLD mutagenesis BP insertion |
| Ymu1_BPinsertion_rev | atcagcagttctcttAGGAGAAGAAGCCATCTC | KLD mutagenesis BP insertion |
| Ymu1_H4Y1_fwd | gaagctgcagtataaagcctatg | KLD mutagenesis H4Y |
| Ymu1_H4Y1_rev | caagaaacctttctctttttcttag | KLD mutagenesis H4Y |
| Ymu1_V304R2_fwd | gacgagccttcgatcgaaactgc | KLD mutagenesis V304R |
| Ymu1_V304R2_rev | catgatacatcagagatgctttttg | KLD mutagenesis V304R |
| Ymu1_K76A1_fwd | tgaagtggatgcgttctcgctccag | KLD mutagenesis K76A |
| Ymu1_K76A1_rev | caaagccatactgtttccg | KLD mutagenesis K76A |
| PjPDS_Ymu1gRNA1_top | TCAACTCGACGAAAAAGCAC | PjPDSg1 cloning |
| PjPDS_Ymu1gRNA1_bottom | GGCCGTGCTTTTTCGTCGAG | PjPDSg1 cloning |
| PjPDS_Ymu1gRNA2_top | TCAATTGCTGTAGTTGGGGC | PjPDSg2 cloning |
| PjPDS_Ymu1gRNA2_bottom | GGCCGCCCCAACTACAGCAA | PjPDSg2 cloning |
| PjPDS_Ymu1gRNA3_top | TCAATTTCTCCGGCCAAGTT | PjPDSg3 cloning |
| PjPDS_Ymu1gRNA3_bottom | GGCCAACTTGGCCGGAGAAA | PjPDSg3 cloning |
| PjDVR_Ymu1gRNA1_top | TCAATGGGTGGATGAAAATG | PjDVRg1 cloning |
| PjDVR_Ymu1gRNA1_bottom | GGCCCATTTTCATCCACCCA | PjDVRg1 cloning |
| PjDVR_Ymu1gRNA2_top | TCAATTGTTGGTTAAGGTGT | PjDVRg2 cloning |
| PjDVR_Ymu1gRNA2_bottom | GGCCACACCTTAACCAACAA | PjDVRg2 cloning |
| PjPDS_Ymu1K76A-gRNA1_top | TCAAAGGGGAAGGTAAGATT | $^{K76A}$PjPDSg1 cloning |
| PjPDS_Ymu1K76A-gRNA1_bottom | GGCCAATCTTACCTTCCCCT | $^{K76A}$PjPDSg1 cloning |
| PjPDS_Ymu1K76A-gRNA2_top | TCAACATCCAGTCTTTAACC | $^{K76A}$PjPDSg2 cloning |
| PjPDS_Ymu1K76A-gRNA2_bottom | GGCCGGTTAAAGACTGGATG | $^{K76A}$PjPDSg2 cloning |
| PjPDS_Ymu1K76A-gRNA3_top | TCAAGAAGTTGAGTGCTTTT | $^{K76A}$PjPDSg3 cloning |
| PjPDS_Ymu1K76A-gRNA3_bottom | GGCCAAAAGCACTCAACTTC | $^{K76A}$PjPDSg3 cloning |
| PjPDSEx6-8for | CTGCAGCCAATGACTCGTTTGT | PDSg2 for native PAGE |
| PjPDS_ISDra2gRNA3_rev | TCAATTTCTCCGGCCAAGTTAGCA | PDSg2 for native PAGE |
| PjPDSEx6-8for | CTGCAGCCAATGACTCGTTTGT | $1^{st}$ PCR PDSg2,3, $^{K76A}$PDSg1-3 |
| PjPDSEx6-8rev | AGAGAGAGTCGTACCTGCAGGA | $1^{st}$ PCR PDSg2,3, $^{K76A}$PDSg1-3 |
| PjDVR_guide1genotype_for | AAGGGTTTTCACATCTCAATTC | $1^{st}$ PCR DVRg1 |

| **Primer name** | **Sequence** | **purpose** |
|---|---|---|
| PjDVR_guide1genotype_rev | GTTAACATTCCTAGGGTTTCGT | $1^{st}$ PCR DVRg1 |
| PjDVR_guide2genotype_for | TTTTCACGTACAGCATTGTCAG | $1^{st}$ PCR DVRg2 |
| PjDVR_guide2genotype_rev | AAACTCCTCTAACGTGTCCTCA | $1^{st}$ PCR DVRg2 |
| PjPDSgenomicfor | GTTGCGAAATCTTGAATTCGGATGA | $1^{st}$ PCR PDSg1 |
| PjPDSgenomicrev | ATCGGTTTATGGCCCGCATC | $1^{st}$ PCR PDSg1 |
| PDSfor1 | atgattcagcggcctatagca**TGATCACG**TGAATTCGGATGATGGCCCAAT | $2^{nd}$ PCR PDSg1 |
| PDSfor2 | atgattcagcggcctatagca**GGTGATGA**TGAATTCGGATGATGGCCCAAT | $2^{nd}$ PCR PDSg1 |
| PDSfor3 | atgattcagcggcctatagca**AACCTACG**TGAATTCGGATGATGGCCCAAT | $2^{nd}$ PCR PDSg1 |
| PjDVR_guide1Amplicon_for1 | atgattcagcggcctatagca**TGATCACG**AAGGGTTTTCACATCTCAATTC | $2^{nd}$ PCR DVRg1 |
| PjDVR_guide1Amplicon_rev1 | cacatcggcgcatagttacg**TTGCTTGC**GTTAACATTCCTAGGGTTTCGT | $2^{nd}$ PCR DVRg1 |
| PjDVR_guide2Amplicon_for2 | atgattcagcggcctatagca**AGGTGATG**ATTTTCACGTACAGCATTGTCAG | $2^{nd}$ PCR DVRg2 |
| PjDVR_guide2Amplicon_rev2 | cacatcggcgcatagttacg**ACTACGGA**AAACTCCTCTAACGTGTCCTCA | $2^{nd}$ PCR DVRg2 |
| PjPDSEx2_rev1 | cacatcggcgcatagttacg**TTGCTTGC**GCATACTTTTAAAATGTTTGCGTCT | $2^{nd}$ PCR PDSg1 |
| PjPDSEx6-7_for1 | atgattcagcggcctatagca**TGATCACG**CTGCAGCCAATGACTCGTTTGT | $2^{nd}$ PCR PDSg2, PDSg3, $^{K76A}$PjPDSg1 |
| PjPDSEx6-7_for2 | atgattcagcggcctatagca**GGTGATGA**CTGCAGCCAATGACTCGTTTGT | $2^{nd}$ PCR PDSg2, PDSg3, $^{K76A}$PjPDSg1 |
| PjPDSEx6-7_for3 | atgattcagcggcctatagca**AACCTACG**CTGCAGCCAATGACTCGTTTGT | $2^{nd}$ PCR PDSg2, PDSg3, $^{K76A}$PjPDSg1 |
| PjPDSEx6-7_rev1 | cacatcggcgcatagttacg**TTGCTTGC**ACCGTTATTCCATCTTGGGCCT | $2^{nd}$ PCR PDSg2, PDSg3, $^{K76A}$PjPDSg1 |
| PjPDSEx6-7_rev2 | cacatcggcgcatagttacg**ACTACGGA**ACCGTTATTCCATCTTGGGCCT | $2^{nd}$ PCR PDSg2, PDSg3, $^{K76A}$PjPDSg1 |
| PjPDSEx7for2 | atgattcagcggcctatagca**GGTGATGA**CACGTGCTTTAATAAAATGAGG | $2^{nd}$ PCR K76A_PjPDSg2, $^{K76A}$PjPDSg3 |
| PjPDSEx7for3 | atgattcagcggcctatagca**AACCTACG**CACGTGCTTTAATAAAATGAGG | $2^{nd}$ PCR K76A_PjPDSg2, $^{K76A}$PjPDSg3 |
| PjPDSEx8rev1 | cacatcggcgcatagttacg**TTGCTTGC**AGAGAGAGTCGTACCTGCAGGA | $2^{nd}$ PCR K76A_PjPDSg2, $^{K76A}$PjPDSg3 |
| PjPDSEx8rev2 | cacatcggcgcatagttacg**ACTACGGA**AGAGAGAGTCGTACCTGCAGGA | $2^{nd}$ PCR K76A_PjPDSg2, $^{K76A}$PjPDSg3 |
| AmpliconEurofins_for1 | ACACTCTTTCCCTACACGACGCTCTTCCGATCT**GTCATGAG**atgattcagcggcctatagca | 3rd PCR, All amplicons |
| AmpliconEurofins_for2 | ACACTCTTTCCCTACACGACGCTCTTCCGATCT**TGCAGCTA**atgattcagcggcctatagca | 3rd PCR, All amplicons |
| AmpliconEurofins_for3 | ACACTCTTTCCCTACACGACGCTCTTCCGATCT**AAGCGACT**atgattcagcggcctatagca | 3rd PCR, All amplicons |
| AmpliconEurofins_for4 | ACACTCTTTCCCTACACGACGCTCTTCCGATCT**GGATATGC**atgattcagcggcctatagca | 3rd PCR, All amplicons |
| AmpliconEurofins_rev1 | GACTGGAGTTCAGACGTGTGCTCTTCCGATCT**GAGAGGTT**cacatcggcgcatagttacg | 3rd PCR, All amplicons |
| AmpliconEurofins_rev2 | GACTGGAGTTCAGACGTGTGCTCTTCCGATCT**GGTAAGCT**cacatcggcgcatagttacg | 3rd PCR, All amplicons |

| Primer name | Sequence | purpose |
|---|---|---|
| AmpliconEurofins_rev3 | GACTGGAGTTCAGACGTGTGCTCTTCCGATCT**CGGAACAA**cacatcggcgcatagttacg | 3rd PCR, All amplicons |
| AmpliconEurofins_rev4 | GACTGGAGTTCAGACGTGTGCTCTTCCGATCT**TGTGGCAT**cacatcggcgcatagttacg | 3rd PCR, All amplicons |

Note: Barcode sequences are shown in bold
